# Video rate volumetric Ca^2+^ imaging across cortical layers using Seeded Iterative Demixing (SID) microscopy

**DOI:** 10.1101/155572

**Authors:** Tobias Nöbauer, Oliver Skocek, Alejandro J. Pernía-Andrade, Lukas Weilguny, Francisca Martinez Traub, Maxim I. Molodtsov, Alipasha Vaziri

## Abstract

Light-field microscopy (LFM) is a scalable approach for volumetric Ca^2+^ imaging with the highest volumetric acquisition rates (up to 100 Hz). While this has enabled high-speed whole-brain Ca^2+^ imaging in small semi-transparent specimen, tissue scattering has limited its application in the rodent brain. Here we introduce Seeded Iterative Demixing (SID), a computational source extraction technique that extends LFM to the scattering mammalian cortex. Using GCaMP-expressing mice we demonstrate SID’s ability to capture neuronal dynamics *in vivo* within a volume of 900×900×260μm located as deep as 380 μm in the mouse cortex and hippocampus at 30 Hz volume rate while faithfully discriminating signals from neurons as close as 20 μm, at three orders of magnitude reduced computational cost. The simplicity and scalability of LFM, coupled with the performance of SID opens up a range of new applications including closed-loop experiments and is expected to propel its wide dissemination within the neuroscience community.

---

Understanding multi-scale integration of sensory inputs and the emergence of complex behavior from global dynamics of large neuronal populations is a fundamental problem in current neuroscience. Only recently, the combination of genetically encoded Calcium (Ca^2+^) indicators (GECIs)^1^ and new optical imaging techniques have enabled recording of neuronal population activity from entire nervous systems of small model organisms, such as *C. elegans*^2,3^ and zebrafish larvae^4,5^, at high speed and single-cell resolution. However, single-cell resolution functional imaging of large volumes at high speed and great depth in scattering tissue, such as the mammalian neocortex, has proven more challenging.

A major limitation is the fundamental trade-off between serial and parallel acquisition schemes. Serial acquisition approaches such as two-photon scanning microscopy (2PM)^6^, provide robustness to scattering, however this is achieved at the expense of temporal resolution. More recently, a number of approaches have been developed to alleviate this restriction^7^ at the cost of increased complexity – by scanning faster using acousto-optic deflectors^8^, remote focusing using mechanical actuators^9^ or acousto-optical lenses^10^, temporal or spatial multiplexing^11–13^, using holographic approaches^14,15^, by selectively addressing known source positions by random access scanning^16–18^, by sculpting the microscope’s point spread function (PSF) in combination with a more efficient excitation scheme^19^ or other PSF engineering approaches^20,21^.

In contrast, parallel or partially parallel acquisition schemes such as wide-field epifluorescence microscopy, different variants of light-sheet microscopy^22,23,5,24,25^, wide-field temporal focusing^2^ and other approaches^26^ can greatly improve temporal resolution. Typically, however, light scattering mixes fluorescence signals originating from distinct neurons and degrades information on their locations when a 2D array detector is used. Thus, parallel acquisition schemes have been mostly limited to highly transparent specimen or to the most superficial regions of scattering tissues such as the mammalian cortex.

Among the parallel acquisition techniques, Light Field Microscopy (LFM)^4,27–30^ has recently been established as a simple yet powerful approach to high-speed, volumetric Ca^2+^ imaging in small semi-transparent model systems such as *C. elegans* and zebrafish larvae^4^. LFM is realized by placing a microlens array in the image plane of a microscope, and a camera in the focal plane of the microlens array. This scheme stands out from competing imaging methods by its unique scalability, as it does not require any time-consuming scanning of the excitation beam to collect 3D information. This effectively decouples imaging speed and field of view (FOV) and thus allows for scaling up volume acquisition rates (limited only by GECI response dynamics and camera frame rate) and FOV at the same time. Moreover, LFM does not require expensive and complex ultrafast laser systems and is not prone to sample heating and nonlinear photo-damage.

While the performance of LFM in transparent samples has been well documented, due to the vectorial and redundant nature of the information it collects^30,31^, LFM also holds the promise of great robustness in scattering samples. However, conventional frame-by-frame reconstruction of LFM images^4,28^ largely fails at harvesting this potential besides being highly computationally resource-intensive.

Here, we introduce an iterative source extraction procedure for scattered LFM data, termed SID (Seeded Iterative Demixing), which achieves accurate neuron localization and signal demixing by seeding inference with information obtained from remnant ballistic light. We iteratively update our estimates of the time series and the scattered images of each active neuron by non-negative, constrained least-squares optimization.

In doing so, we achieve robust signal demixing up to a depth of about four scattering lengths, corresponding to up to ~400 μm in mouse cortex, and demonstrate an increased temporal and spatial fidelity when SID is applied to weakly scattering samples, such as larval zebrafish. We demonstrate acquisition of neuronal activity at 30 Hz within densely labeled volumes of ~900 × 900 × 260 μm at depths up to ~380 μm in mouse cortex and hippocampus. Finally, we show that SID provides three orders of magnitude of improvement in computational resource requirements, compared to conventional LFM reconstruction and post-processing. It thus opens the door to a range of qualitatively new applications, including real-time whole-brain recording, and closed loop interrogation of neuronal population activity in combination with optogenetics and behavior.

## RESULTS

### Seeded Iterative Demixing (SID) light field microscopy demixes neuronal signals in a scattering medium

On average only about 34% of incident photons still travel in their original direction (“ballistic photons”) after propagating the characteristic distance of one scattering length (~50-100 μm for visible light in the cortex^32^), while the remaining photons experience scattering that deflects them from their original direction of propagation. In brain tissue, scattering is strongly peaked in the forward direction, so that some directional information is preserved even after several scattering events^32^.

In conventional wide-field fluorescence imaging, scattering causes image features to appear blurred and overlapping, rendering their segmentation a highly ill-posed mathematical problem. In contrast to single-photon techniques^5,25,26^, including the various implementations of light sheet microscopy, LFM allows for resolving directional information retained in the scattered light field, and hence facilitates the extraction of neuronal signals from the raw LFM data (Supplementary Note 1 and Supplementary Fig. 1).

Our SID method effectively achieves this by (1) exploiting the high-resolution spatial information contained in remnant ballistic light, (2) incorporating knowledge on the LFM PSF and the effects of scattering, and (3) employing a constrained spatiotemporal matrix factorization technique to demix the effects of scattering by utilizing both the spatial and temporal information present in the data, without any assumptions on source positions or signal characteristics (Fig. 1a, Methods, Supplementary Note 2, Supplementary Fig. 2).

**Figure 1.**
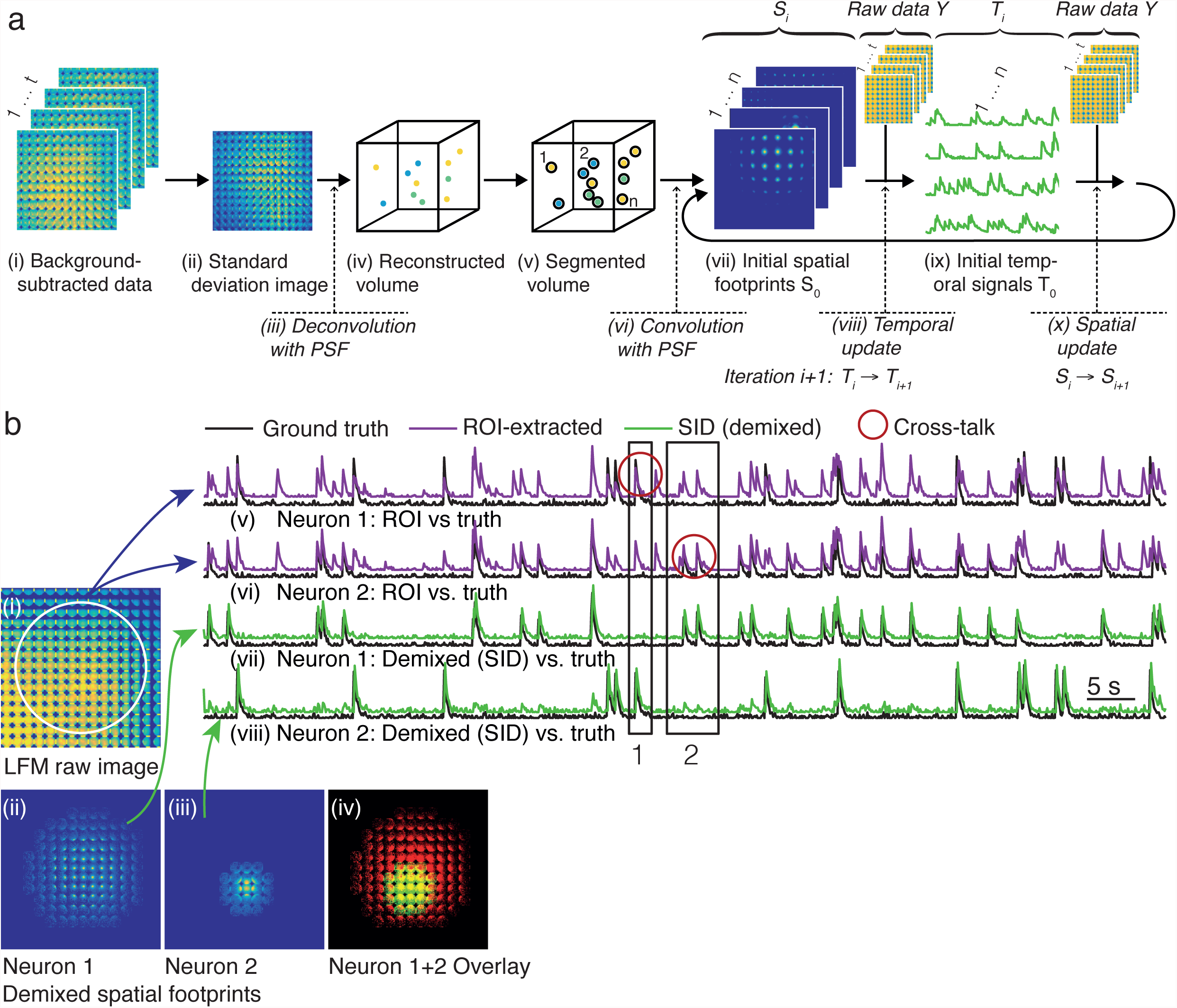
Seeded iterative demixing of light-field recordings in scattering tissue. **(a)** Illustration of key steps in the Seeded Iterative Demixing (SID) algorithm (see Methods): After background subtraction and detrending, a standard deviation image is calculated. This allows to extract the remaining unscattered light from active neurons since unscattered photons are spread onto fewer pixels. The standard deviation image is reconstructed as a single image by deconvolution with a numerically simulated PSF. The resulting volume is segmented to obtain an initial guess of neuron positions from which a set of neuronal footprints on the sensor is generated. The corresponding initial temporal signals, the subsequent updated spatial footprints and temporal signals are found by iteratively solving a bi-convex, non-negative constrained matrix factorization problem. **(b)** Demixing of overlapping LFM neuron footprints by Seeded Iterative Demixing (SID). (i) Standard deviation image of the synthetic LFM raw movie. (ii) and (iii) two demixed spatial components (neuron footprints) detected within the encircled region shown in (i). (iv) Composite color overlay of the spatial components (ii) and (iii), showing considerable overlap (yellow). (v-viii) Comparisons of ground truth temporal signals extracted using SID and spatial region-of-interest extraction. Black: Ground truth signals for the two neurons whose LFM images fall into the area encircled in (i). Green: demixed signals corresponding to the spatial components (ii) and (iii), as indicated by the arrows. Violet: signal extracted by summing over the encircled area in (i). Black rectangles and red circles highlight intervals where time series of neuron 1 (ii) and neuron 2 (iii) are mixed for a circular region-of-interest extraction (v, vi) and demixed for SID (vii, viii) as evident from comparison of the corresponding signals to the ground truth.

To verify and characterize the demixing performance of our SID approach, we first applied it to synthetic datasets containing simulated scattering tissue and randomly positioned neurons with partially correlated, GECI-like activity (see Supplementary Note 3). Application of our SID algorithm to synthesized data corresponding to a depth of ~400 μm in mouse cortex allowed us to reliably demix overlapping spatial footprints (Fig. 1b). In cases where signal extraction based on regions-of-interest (ROI) would give highly mixed signals (Fig. 1b, violet traces), SID allowed for faithful signal demixing, yielding close correspondence (mean correlation of 0.76) of the extracted signals (Fig. 1b, green traces) with the ground truth signal (black traces). We find that SID requires only a small difference in temporal activity and spatial footprint to faithfully differentiate the two entities. As a result, laterally overlapping neurons in slightly different axial planes can be resolved by SID (see Supplementary Fig. 3).

### Seeded Iterative Demixing (SID) improves source localization in zebrafish larvae

Zebrafish larvae are an ideal testbed for establishing a baseline for *in vivo* performance of our SID approach in the weakly scattering regime, while the higher neuron density in the larval zebrafish brain compared to the mammalian cortex poses an additional challenge to our method and requires a method with higher signal discriminability.

We built a hybrid 2PM-SID microscope (Fig. 2a, and Methods) and compared the neuron positions extracted via SID to a high-resolution 2PM image stack, within a volume of 775 × 195 × 200 μm in the anterior part of the zebrafish spinal cord (Fig. 2b). The spatial resolution of our LFM, based on standard reconstruction, was 3.5 μm laterally, corresponding to about half a neuron diameter in zebrafish larvae^5^ and 9 μm axially. Spatial segmentation of the 2PM stack yielded a total of 1337 active as well as inactive neurons within the above volume. (Fig. 2b shows a shorter depth range for clarity). SID inherently detects active neurons only, and yielded 508 neurons whose positions clearly coincide with neurons in the 2PM stack. We then recorded spontaneous neuronal activity from the entire larval zebrafish brain covering a volume of 700 × 700 × 200 μm at 20 Hz for 240 seconds. In this case SID found a total of 5505 active neurons, the positions in 3D and the maximum intensity projection of which are shown in Fig. 2c-e (for corresponding activity time series see Supplementary Fig. 4). We compared and assessed the validity of these results by analyzing the same data using conventional reconstruction, followed by Independent Component Analysis (ICA)^33^, as used previously^4^. We found a good overall agreement, while showing SID’s robustness to crosstalk in contrast to ICA, which is known to suffer from crosstalk when applied to highly correlated signals^34^ (see Supplementary Note 4 and Supplementary Fig. 5 for detailed results and a discussion of other extraction methods).

**Figure 2.**
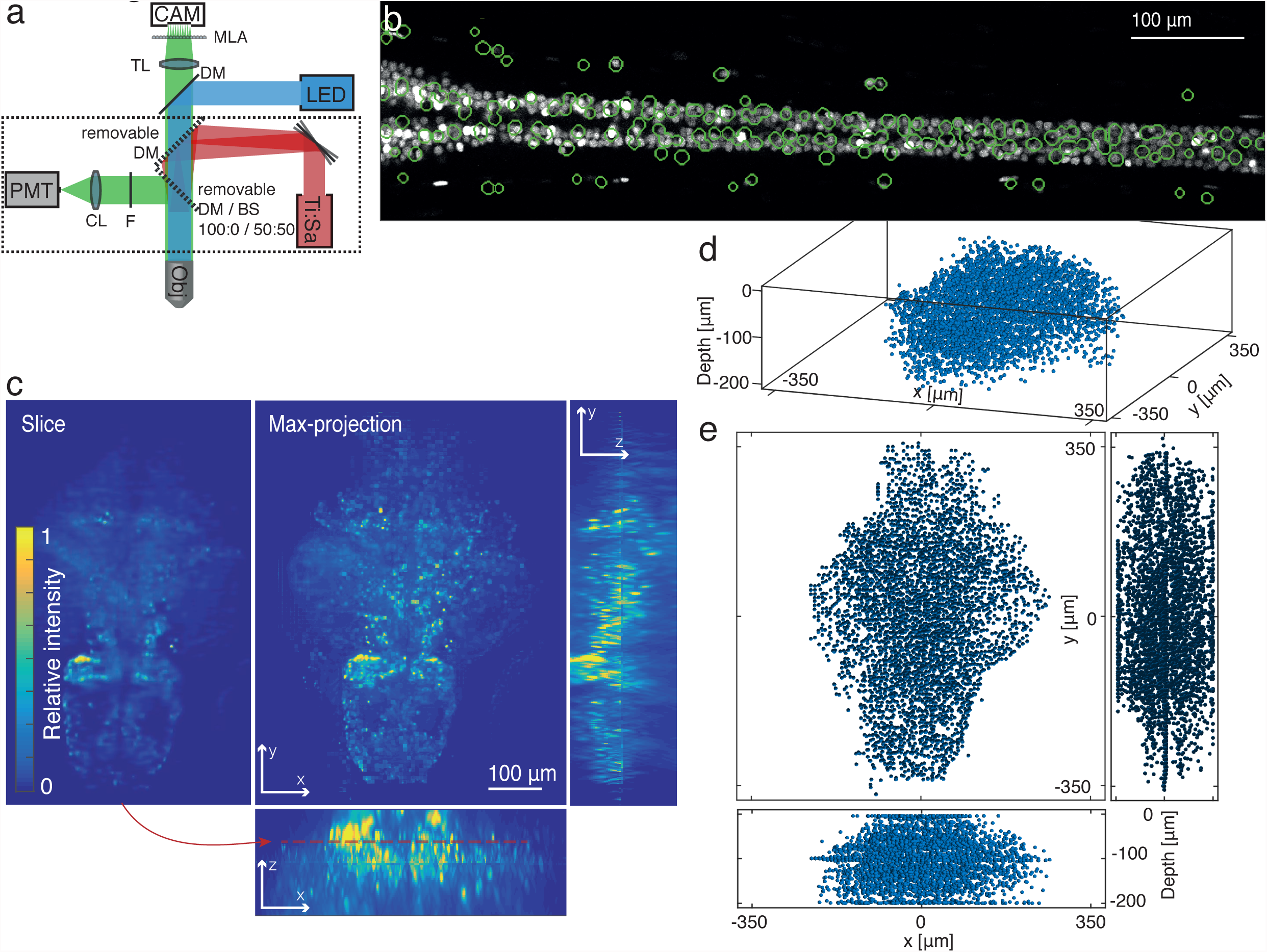
Seeded iterative demixing applied to whole-brain Ca^2+^ imaging in larval zebrafish. **(a)** Schematic of the hybrid LFM and 2PM experimental setup, as described in the Methods. LED, light emitting diode light source. Ti:Sa, Titanium:Sapphire laser. DM, dichroic mirror. BS, beam splitter. F, emission filter. CL, collection lens. TL, tube lens. MLA, microlens array. CAM, sCMOS camera, PMT, photomultiplier tube. **(b)** 2PM image of an anterior part of the spinal cord in larval zebrafish labelled with nucleus-confined GCaMP6s, overlaid with outlines of active neurons (green circles) detected using Seeded Iterative Demixing (SID) in a five minute LFM movie (5 fps) of the same location. **(c)** Frontal slice and maximum-intensity projections along three orthogonal directions of the reconstructed LFM standard-deviation image, same recording as in (d-e). Obtained using total-variation and sparsity constraints during reconstruction for artefact reduction and optimized contrast. This data was used as input to the SID segmentation step to seed spatio-temporal SID demixing. **(d)** Isometric rendering of the neuron locations detected by Seeded Iterative Demixing (SID), corresponding to the neuronal signals shown in Supplementary Fig. 4. **(e)** Orthogonal views onto the neuron locations shown in (d).

### Seeded Iterative Demixing (SID) enables high-speed volumetric Ca^2+^ imaging in mouse cortex and hippocampus at 380 μm depth

The severity of degradation of image and neuronal signal quality due to scattering and reconstruction artifacts in standard LFM reconstruction becomes strikingly apparent when *in vivo* data recorded in the posterior parietal cortex (PPC) of awake mice is conventionally reconstructed (Fig. 3a-b, Supplementary Video 1 and 2). When applying SID to the same data at various depths, the effectiveness of our method becomes evident. Using SID, we could resolve individual neurons in 3D and their corresponding signals within a FOV of ~900 μm diameter up to a depth of 380 μm at a volume acquisition rate of 30 Hz with high fidelity (see Fig. 3c-d for neuron positions, Supplementary Fig. 6 for activity time series, and Supplementary Video 3 and 4). Thereby we could capture locations and activities of neurons in mouse cortical layers I-III and part of the layer IV at 30 Hz volume rate with only two successive recordings. Our algorithm identified over 500 active neurons during a 60 second recording (Supplementary Fig. 6), corresponding to ~10% of all labeled neurons (5296) as identified using a high-resolution 2PM stack. Of the total number of identified neurons, 296 were in the depth range from zero to 170 μm, and 208 active neurons in the range from 120 to 380 μm.

**Figure 3.**
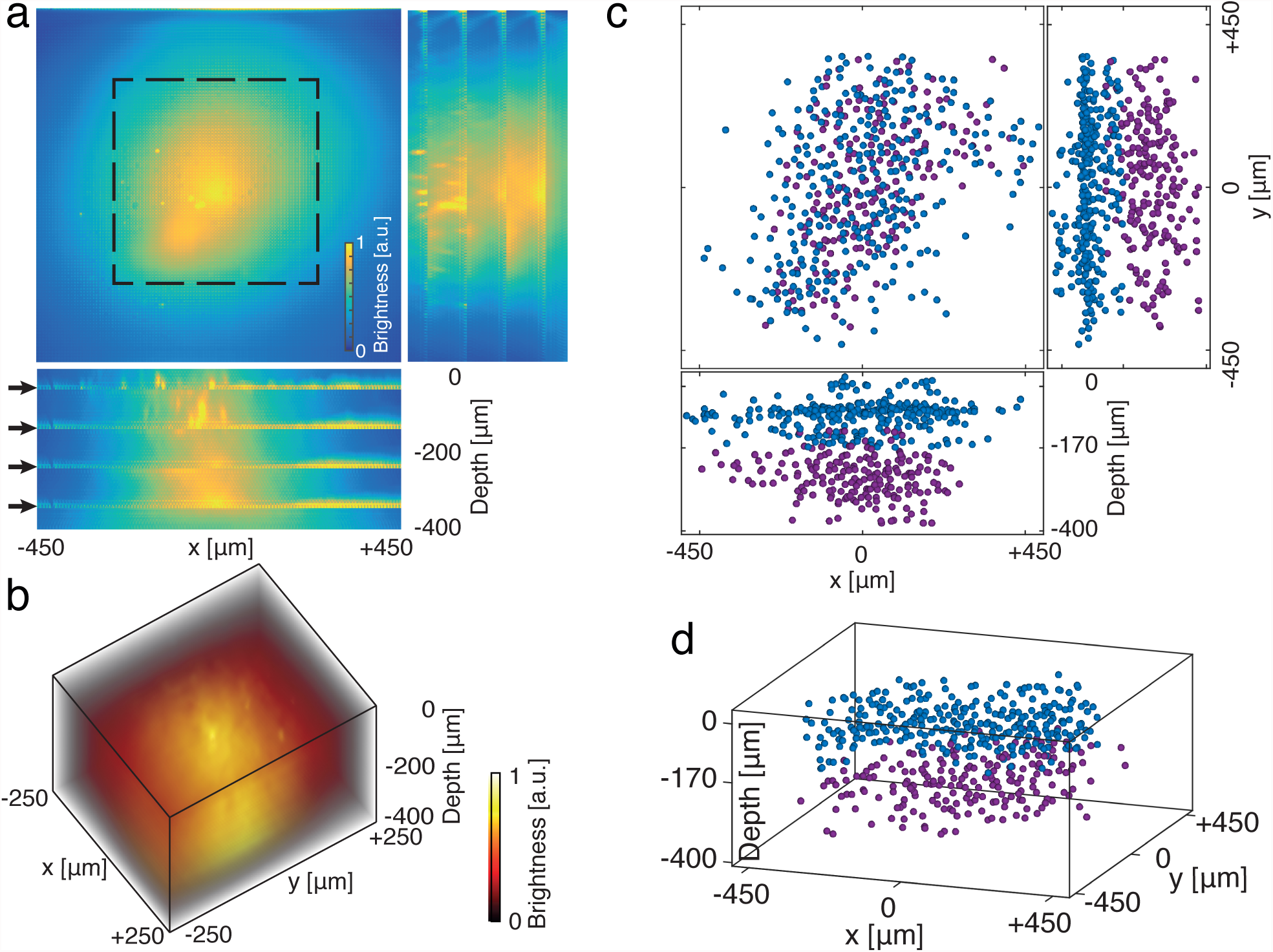
Video-rate volumetric Ca^2+^ imaging to 380 μm depth in mouse cortex. **(a)** Maximum-intensity projections of a stack of volumetric frames obtained by conventional reconstruction of LFM recordings from the mouse cortex. Five independent recordings at increasing depths are shown. Reconstruction artefacts in classical LFM reconstruction are visible in the side views as high-intensity planes (indicated by arrows). Dashed rectangle indicates lateral region shown in (b). **(b)** Rendering of isometric perspective onto the data shown in (a), zoomed into lateral region indicated by dashed rectangle in (a). **(c)** Neuron positions obtained by Seeded Iterative Demixing (SID) from a one-minute recording at 30 fps in mouse cortex. Neuron positions from two subsequent recordings of different depth ranges are shown together, color indicates recording range: 0-170 μm (blue) and 120-380 μm (red). **(d)** Isometric perspective onto data shown in (c).

To further illustrate the versatility of SID, we applied our method to imaging of CA1 hippocampal neurons using a cranial window implanted after cortical aspiration^35,36^ (Fig. 4a). Capturing the neuronal population activity within a volume of ~900 × 900 × 200 μm containing the cell body layer of CA1 neurons, we could reliably identify, extract and demix Ca^2+^ signals from 150 neurons arranged in the curved layer geometry typical of the anatomy of this region (Fig. 4b-d). Example neuron traces showed robust and pronounced Ca^2+^ transients that are consistent with the high-frequency bursts of neuron types in this brain region^37^ (Fig. 4e).

**Figure 4.**
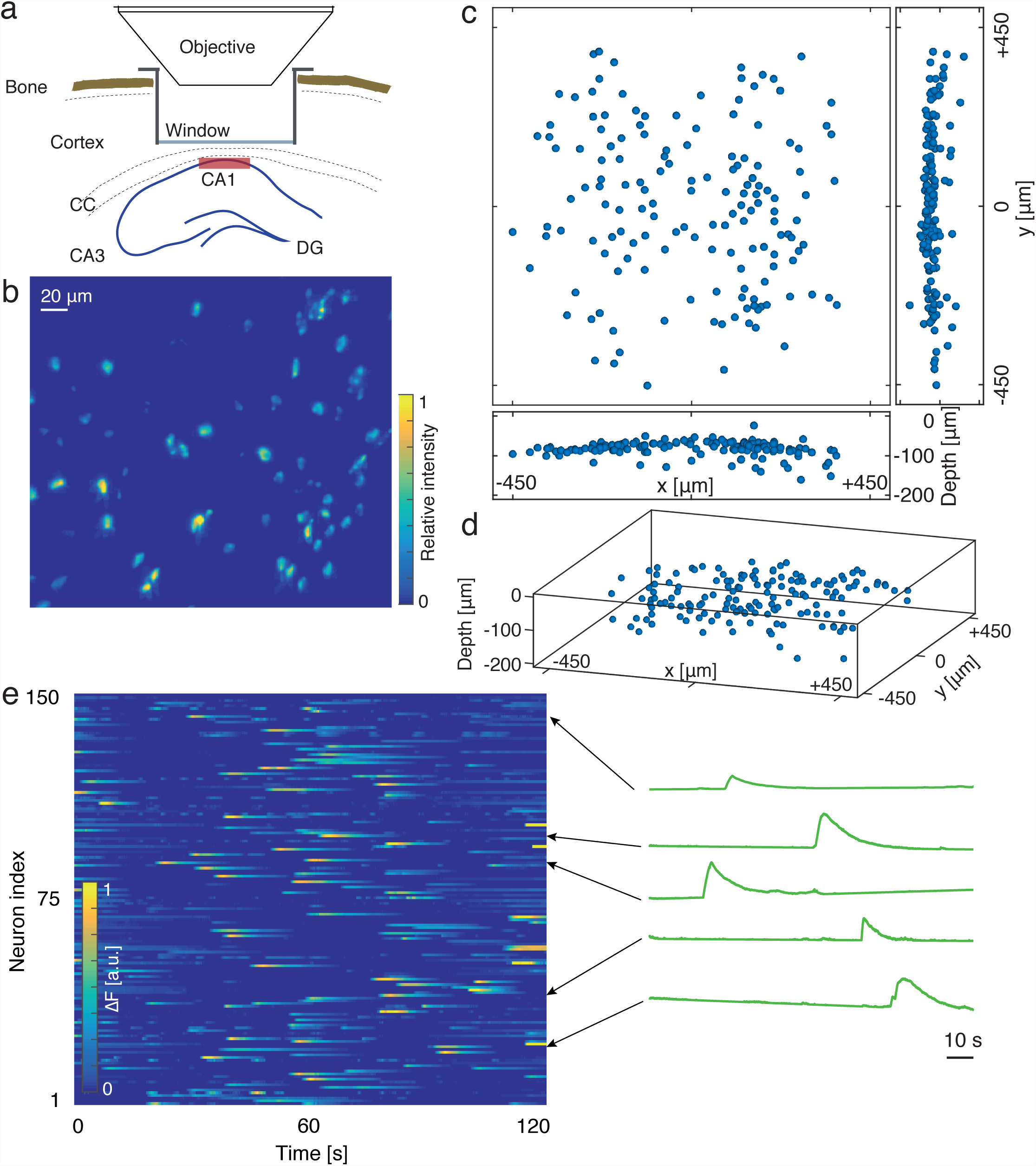
Video-rate volumetric Ca^2+^ imaging in mouse hippocampus. **(a)** Schematics of the hippocampal window preparation, indicating corpus callosum (CC), and region of hippocampus proper *Cornu Ammonis* (CA1, CA3) and dentate gyrus (DG). The red box indicates the approximate imaging volume. **(b)** Maximum-intensity projection along optical axis of reconstructed standard deviation LFM image, showing GCaMP6m-labelled neurons active during a two-minute, 30 fps recording. **(c)** Positions of 150 active, GCaMP6m-labelled neurons obtained by Seeded Iterative Demixing (SID) of a two-minute, 30 fps LFM recording in mouse CA1. **(d)** Isometric perspective onto data shown in (d). **(e)** Heatmap of neuron activity traces obtained by Seeded Iterative Demixing (SID), corresponding to positions shown in (c) and (d). Right panel: Four representative neuron traces as line plots, with their positions in the heatmap indicated by arrows.

### Seeded Iterative Demixing (SID) reduces computational cost by three orders of magnitude

An key feature of our method is that it allows to extract signals and source positions from LFM recordings without the enormous computational resources required by previous frame-by-frame deconvolution approaches^4,28^ and ICA analysis. SID decreases the computation time by a factor of approximately 1,000 for a typical example of a 10,000-frame recording (Fig. 5a). Reconstructing such a recording requires ~8,000 GPU-hours (GPU - graphics processing unit) on a high-end GPU cluster when using the classical frame-by-frame reconstruction approach^4^. In addition, ICA analysis of a reconstructed dataset requires computational resources beyond what is typically available on a workstation (Supplementary Note 4). In contrast, SID requires the full reconstruction of only one frame out of the 10,000 frames, reducing the procedure to approx. 7 GPU-hours and making it suitable for execution on a single workstation.

**Figure 5.**
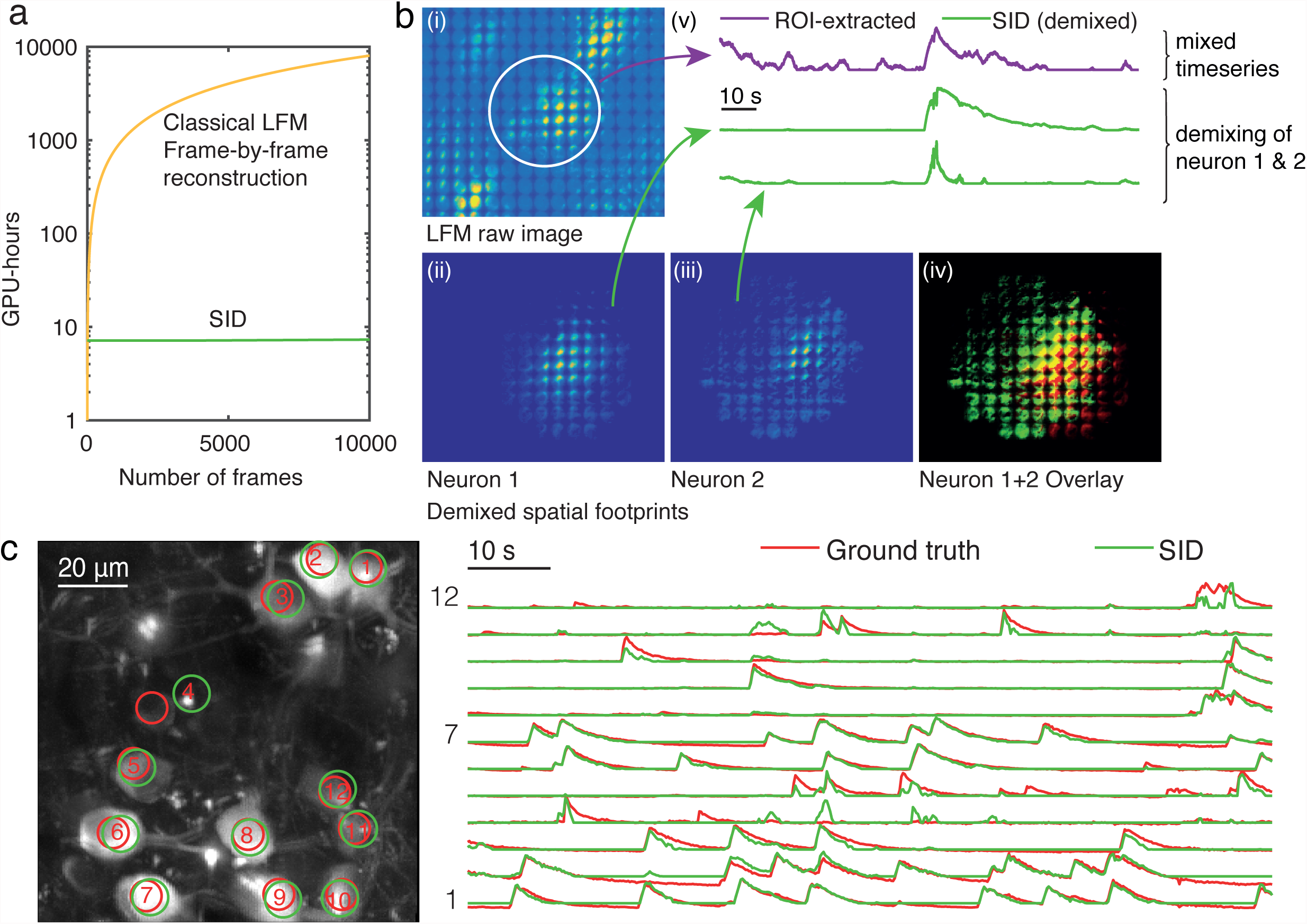
Experimental verification of Seeded Iterative Demixing (SID) performance and computational cost. **(a)** Computational cost of conventional reconstruction (orange) and Seeded Iterative Demixing (SID) (green) of LFM recordings vs. number of frames. Vertical axis gives approximate computation time required on a nVidia GTX Titan Black GPU using a Matlab implementation. Note the logarithmic vertical axis. **(b)** Demixing of overlapping LFM neuron footprints by Seeded Iterative Demixing (SID) in hippocampal recordings. (i) Region from the standard deviation image of LFM raw movie. (ii) and (iii) two demixed spatial components (neuron footprints) detected within the encircled region shown in (i). (iv) Composite color overlay of the spatial components (ii) (green) and (iii) (red), showing considerable overlap (yellow). (v) Green: demixed signals corresponding to the spatial components (ii) and (iii). Violet: signal extracted by summing over the encircled area in (i). **(c)** Comparison of neuron positions and activity traces as obtained from segmentation of a planar 2PM recording (red circles and traces), and Seeded Iterative Demixing (SID) (green circles and traces) of a simultaneous LFM recording of the same region, as described in the main text. Background image in left panel is a maximum projection along time. Numbers 1-12 in image identify corresponding traces in left panel.

### Seeded Iterative Demixing (SID) allows for demixing and localization of overlapping neuronal signals in the mouse brain with time series consistent with 2PM ground truth

Next, we experimentally and systematically demonstrated the capability of SID to demix neuronal signals in scattering tissue while providing neuronal time series that closely match those obtained by more established methods such as 2PM. As an example on the single-neuron level, we selected two CA1 neurons that were indistinguishable based on their spatial senor footprints, and which exhibit highly correlated activity (Fig. 5b). As shown in Fig. 5b, SID can detect them as individual neurons spatially and demix their corresponding time signals. To achieve this, SID only requires a few pixels within the spatial footprint of each neuron to eliminate crosstalk from the respective other neuron (see also Supplementary Fig. 3).

The volumetric FOV and frame rate of our method exceed those of other techniques, such as 2PM, that are typically used for *in vivo* Ca^2+^ imaging at similar depths in the mouse cortex. It is therefore impossible to establish an experimental ground truth for our method by directly comparing the neuronal time series obtained by SID and 2PM within our typical volume sizes and volume acquisition rates. Nevertheless, we generated experimental ground truth data and validated that time series extracted by SID are indeed consistent with data from more established methods such as 2PM, within the limits of current technology. To do so, we used a hybrid 2PM-SID microscope (see Methods). We performed 2PM excitation in a single plane in the mouse cortex while simultaneously detecting the fluorescence using our SID detection arm and a photomultiplier tube (PMT) point detector in our hybrid 2PM-SID (Fig. 2a). Our 2PM hardware allowed us to scan a plane of 200 × 200 μm at 5 Hz. Fig. 5c compares localization and signal extraction for twelve neurons found in this region using spatial segmentation based on watershed transform on the obtained 2PM data (red circles), and SID on data obtained in the LFM detection arm (green circles). It clearly demonstrates that signals extracted by SID are in quantitative agreement with 2PM recordings (12 out of 12 active neurons detected; mean cross-correlation of signals from the two methods: 0.85).

To obtain a more comprehensive and quantitative evaluation of the performance of SID, we recorded a set of single-plane, simultaneous 2PM-SID movies at a series of axial depths (100–375 μm, total n = 18 recordings). Neuron positions and signals were extracted from the 2PM channel using a recently published and increasingly used method^34^ based on constrained matrix factorization (“CaImAn”). The output of CaImAn was assessed and corrected manually to establish a ground truth, to which we quantitatively compared both the raw CaImAn output and SID (see Supplementary Note 5 for additional details). We found that SID offers a comparable or better compromise between sensitivity (Sensitivity score) and robustness (Precision score), resulting in slightly higher overall performance (F-Score) (Supplementary Note 5 and Supplementary Fig. 7).

The quality of the SID-extracted neuronal activity traces compared to ground truth was characterized at different depths in Fig. 6. We found that the median correlation between SID-extracted and 2PM ground truth signals decays only slightly (Fig. 6a), from 0.92 ± 0.06 at 100 μm depth to 0.90 ± 0.06 at 375 μm (median ± standard error of the median). Of all true positive SID detections, 73% have a correlation with ground truth of better than 0.8, and 60% better than 0.9 (Fig. 6b-c) while only 10% of extracted signals exhibit a low (< 0.4) correlation with 2PM ground truth and correspondingly a degraded overlap of the neuronal signal due to crosstalk with nearby neuropil. To gain an insight into the dependence of such mismatches on tissue depth, we calculated how the fraction of SID-extracted neurons with a correlation to ground truth of less than 0.5 depended on tissue depth (Fig. 6d). We found their fraction to represent only 6% at 100 μm depth and about 12% at 375 μm. While this shows that SID can correctly identify and assign neuronal signals for the vast majority of neurons even in a densely-labeled sample, and since the main source of the above mismatches was interaction with the neuropil, these values can be improved in future. This can be done by eliminating neuropil labelling by using soma- or nucleus-confined Ca^2+^ indicators. In addition, we also outline a computational strategy for demixing and rejecting neuropil contributions from the signals in Supplementary Note 6 and Supplementary Fig. 8.

**Figure 6.**
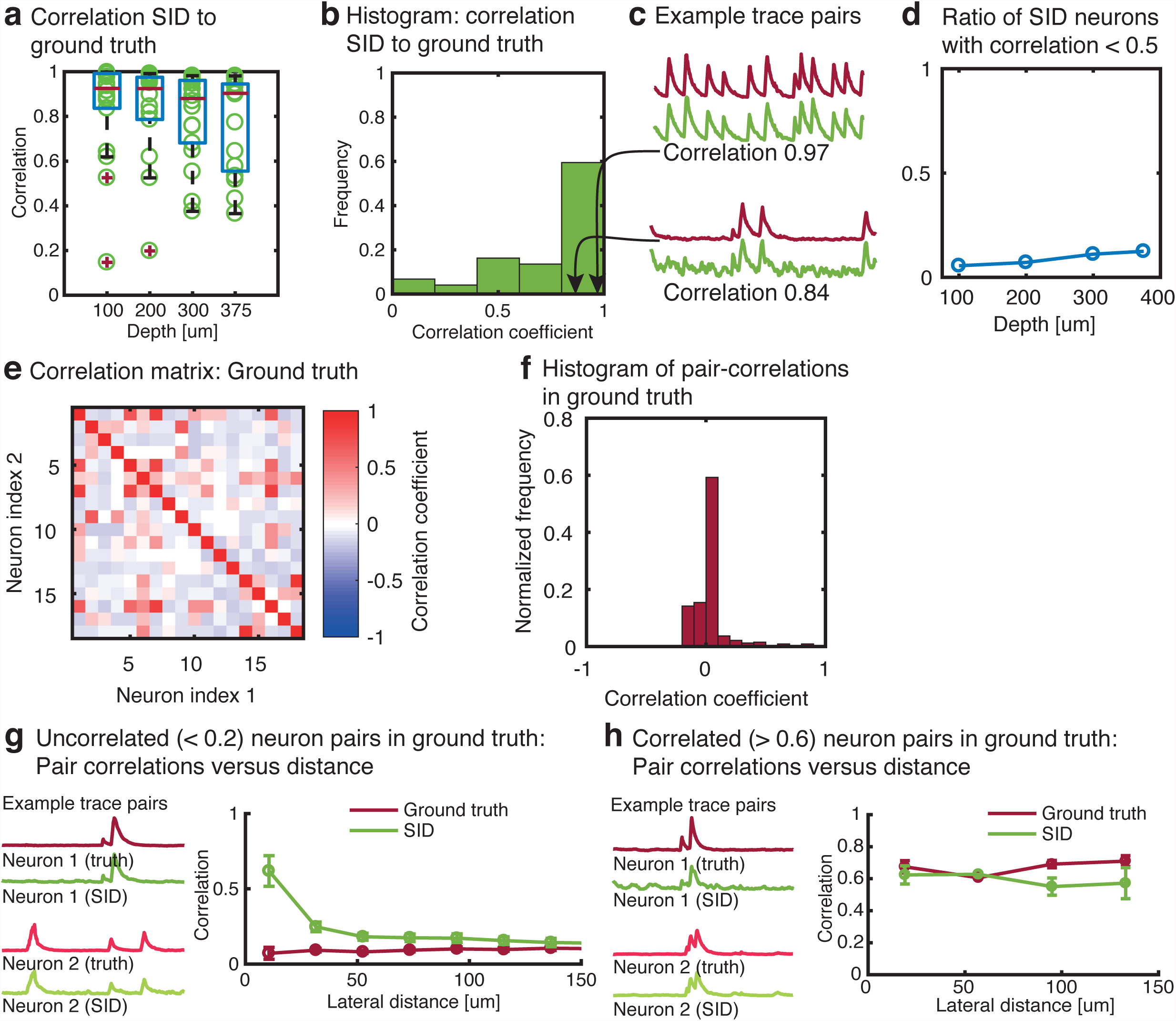
Statistical analysis of Seeded Iterative Demixing (SID) neuron detection and signal extraction performance based on simultaneous 2PM-SID recordings. **(a)** Comparison of SID-extracted signals to ground truth: Correlation versus depth and **(b)** histogram of correlation coefficients of SID signals and their ground truth counterparts, shown for one example recording. Red crosses in (a) indicate box plot outliers, green circles are data points. **(c)** Examples of two SID (green) and corresponding ground truth (red) signal pairs and their respective correlation coefficients. **(d)** Ratio of SID-signals with correlation to ground truth of less than < 0.5 versus imaging depth. **(e)** Comparison of SID performance for pairs of correlated/uncorrelated ground truth neurons: Correlation matrix for ground truth activity signals in one example recording. **(f)** Histogram of the values shown in (e). **(g)** Pair correlations for uncorrelated (< 0.2) neurons extracted by SID and their corresponding ground truth pair correlations (error bars: s.e.m.). Left panel: Example traces for an uncorrelated neuron pair in ground truth, and the corresponding SID neurons. **(h)** Same as in (g), pre-selected for correlated neuron pairs in ground truth (correlation > 0.6). Left panel: Example signals for a correlated neuron pair in ground truth, and the corresponding SID neurons.

Next, we aimed to investigate SID’s performance in demixing signals of nearby neurons. Both physiological correlations of neuronal signals – which are known to generally increase with decreasing neurons pairs distances – as well as degradation of SID’s performance at short neuron pair distances are expected to result in an increase in the observed correlation for decreasing distance of neuron pairs. To dissect the underlying drivers of such observed correlations for the SID-extracted pairs, we investigated their dependence on whether the underlying ground truth pair dynamics was correlated or uncorrelated. To identify such ground truth neuronal pairs, we calculated the corresponding cross-correlation matrix and histogram (Fig. 6e-f). We subsequently selected all uncorrelated neuronal pairs (<0.2) as well as correlated neuronal pairs (>0.6) and examined the correlations of the corresponding signal pairs in SID (Fig. 6g-h). We found an increase in correlation for pairs of uncorrelated ground truth neurons for separations smaller than ~20 μm, while for pairs with correlated ground truth activity the corresponding SID-extracted pairs exhibited a similar correlation as their ground truth pairs over a range of lateral distances, and for as close as ~20 μm. The above un-physiological increase in the observed correlation for uncorrelated ground truth neuron pairs extracted by SID (Fig. 6g) below ~20 μm, as well as the consistency of SID with correlated ground truth pairs down to approximately the same distance, provides us with a metric that represents the discriminability achieved by our SID algorithm, i.e. its ability to detect and assign neuronal time series in the scattering mouse brain.

## DISCUSSION

In this work we have introduced SID, a new, scalable approach for recording volumetric neuronal population activity at high speed and depth in scattering tissue. We have shown that using SID, neuronal activity traces of cells expressing genetically encoded Ca^2+^ indicators can reliably be extracted within a volume of ~900 × 900 × 260 μm at 30 Hz in the mouse cortex, from neurons located as deep as 380 μm and at a discriminability performance of 20 μm, as well as from similarly sized volumes in the mouse hippocampus. Moreover, we show that SID can achieve this performance in the presence of active neuropil. Our method highlights the potential of combining optical imaging with jointly designed computational algorithms to extract information from scattering media, as also demonstrated by other recent developments in signal extraction techniques^38–40^.

Compared to other existing methods for high-speed volumetric Ca^2+^ imaging^8,14,16–21^, SID stands out by its combined acquisition volume and speed, its simplicity and exceptionally low cost as well as its extreme scalability, which we expect to contribute to its rapid dissemination.

Also, unlike in the above techniques, the voxel acquisition rate and resolution in SID are *independent* of the size of the acquired sample volume and only limited by the camera frame rate (up to 100 Hz) and fluorophore properties. It is therefore conceivable to extend SID to much larger FOVs without sacrificing its performance in speed and resolution.

The depth penetration of our method is currently limited by background fluorescence emerging from below the reconstructed volume. This problem can be addressed partially at the expense of increased computational cost, e.g. by modeling PSFs with a larger axial range, which would be able to explain more of the recorded light in terms of localized sources rather than in terms of a diffuse background. Labelled and active neuropil contributes to this background, and hence soma-confined or nucleus-restricted Ca^2+^ reporters could help to increase the obtainable depth range of our method and the quality of the extracted signals. In its current realization however, we show that neuropil signal can be demixed and rejected based on a shape-sensitive approach. Further, several strategies exist in case applications should also require identifying inactive neurons (see Supplementary Note 6).

Beyond this, faithful extraction of neuronal signals will be limited by the loss of directional information due to multiple photon scattering. The critical depth for information loss is known as the transport mean free path and depends on the scattering length anisotropy parameter. In the mouse brain, it amounts to ~10 scattering lengths, or 500-1000 μm^32^.

While previous implementations of LFM required cluster-scale resources, SID enables volumetric signal extraction across multiple cortical areas and layers at unprecedented speed using simple optical components and a workstation-grade computer. Thus, SID represents a transformative step, as the three-order-of magnitude reduction in computational burden demonstrated here opens the door to real-time and closed loop applications, as well as the conception of new LFM-derived volumetric imaging approaches far exceeding existing scale and versatility.

## ACKNOWLEDGMENTS

We thank W. Haubensak and his lab members for sharing animal facility and reagents, M. Colombini and the IMP workshop for manufacturing of mechanical components, F. Schlumm and Q. Lin for help with zebrafish imaging. The computational results presented here have been achieved in part using the Vienna Scientific Cluster (VSC). T.N. acknowledges the Leon Levy Foundation (Leon Levy Fellowship in Neuroscience). This work was supported in part through funding from the Vienna Science and Technology Fund (WWTF) project VRG10-11, the Human Frontiers Science Program Project RGP0041/2012, the Institute of Molecular Pathology, the Kavli Foundation, and the Intelligence Advanced Research Projects Activity (IARPA) via Department of Interior/Interior Business Center (DoI/IBC) contract number D16PC00002. The US Government is authorized to reproduce and distribute reprints for Governmental purposes notwithstanding any copyright annotation thereon. The views and conclusions contained herein are those of the authors and should not be interpreted as necessarily representing the official policies or endorsements, either expressed or implied, of IARPA, DoI/IBC, or the US Government.

## AUTHOR CONTRIBUTIONS

T.N. contributed to the conceptualization of the imaging and signal extraction approach, designed and built the imaging system, performed experiments, wrote software, analyzed data and wrote the manuscript. O.S. contributed to the conceptualization and implementation of the signal extraction approach, wrote software, analyzed data and contributed to the writing of the manuscript. A.J.P.-A. performed virus injections, cranial window surgeries in hippocampus and cortex, performed imaging experiments, and contributed to the writing of the manuscript. F.M.T. and L.W. performed virus injections, cranial window surgeries and performed imaging experiments. M.I.M. contributed to generation of the synthetic datasets and simulations. A.V. conceived and led the project, conceptualized the imaging and signal extraction approach, designed *in vivo* mouse experiments, and wrote the manuscript.

## COMPETING FINANCIAL INTERESTS

The authors declare no competing financial interests.

## METHODS

### Hybrid light field and two-photon microscope

The microscope used for simultaneous 2PM and LFM imaging, the fish recordings and mouse recordings (Fig. 2a) is built around a Scientifica Slicescope platform with a custom LFM detection arm.

The two-photon excitation source (Coherent Chameleon) delivered 140 fs pulses at 80 MHz repetition rate and 920 nm wavelength. The beam intensity was controlled via an electro-optical modulator (Conoptics) for attenuation and blanking, and fed into the a galvo-based scan head (Scientifica). The 2P path and the one-photon excitation/LFM detection path were combined via a short-pass dichroic mirror (Semrock FF746-SDi01). One-photon excitation light from a blue LED (CoolLED pe-2) was fed into an Olympus epi-fluorescence illuminator and reflected into the LFM detection path via a standard EGFP excitation filter and dichroic.

Depending on the experiment, either one-photon or two-photon light was used while the other was blocked. Either was focused by a Nikon 16x 0.8NA water-dipping physiology objective into the sample. For zebrafish experiments, Olympus 20x 1.0NA and Olympus 20x 0.5NA water-dipping objectives were used.

Fluorescence from the sample was detected either by a non-descanned PMT arm, or the LFM arm, or split among both. The split ratio was determined by a main beamsplitter inserted into the beam path behind the objective. A custom detection head design allowed for quick switching between configurations that route 100% to the PMTs (665 nm long-pass dichroic, Scientifica), 100% to the LFM arm (no filter), or split the fluorescence 10:90 or 50:50 (PMT:LFM) (Omega 10% beam sampler or Thorlabs 50:50 vis beam splitter, respectively). The PMT detection arm consisted of an IR blocking filter, collection lens, 565LP dichroic, and 525/50nm and 620/60nm emission filters with, and Scientifica GaAsP (green channel) and alkali (red) PMT modules.

For LFM detection, fluorescence passed through the short-pass dichroic that couples the laser into the beam path, as well as the one-photon filter cube. The image formed by a standard Olympus tube lens was then relayed via two 2-inch achromat lenses (f = 200 mm, Thorlabs) onto a microlens array (MLA, Okotech, custom model, size 1” square, f-number 10, 114 μm microlens pitch, quadratic grid, no gaps). The f-number of the MLA was matched to the output f-number of the microscope. The back focal plane of the MLA was relayed by a photography macro objective (Nikon 105 mm/2.8) at unity magnification onto the sensor of an Andor Zyla 5.5 sCMOS scientific camera, which can be read out at up to 75 Hz at full resolution (2560 × 2160 px, 16 bit).

The setup was controlled from a dual-CPU workstation (HP Z820) with four solid-state disks in a RAID-0 configuration for fast image acquisition and National Instruments 6110 and 6321 cards for analogue and timing I/O. Experiments were controlled using Micro-manager and Scanimage for the one-photon and two-photon parts of the setup, respectively.

### Source extraction algorithm and data analysis

Our source extraction approach (see Supplementary Fig. 2 for illustrations of algorithm steps and Supplementary Note 2 for further discussions) starts with a rank-1 matrix factorization of the time series of raw images to remove background and common-mode dynamics. A motion detection metric (see Supplementary Note 7 and Supplementary Fig. 9) is computed on the background-subtracted images, and frames with a motion metric value above threshold are excluded from further processing. Next, we compute the standard deviation of each pixel along time, resulting in a “standard deviation image”. We deconvolve the standard deviation image using a Richardson-Lucy-type algorithm (with non-negativity and, optionally, sparsity constraints, see Supplementary Note 2) and a numerically simulated PSF, as described previously^4,28^. This results in a volumetric frame containing neurons that are active in the recording as bright regions. The reconstructed volume is band-pass filtered and segmented using a local maximum search, resulting in a dictionary of neuron candidate positions. Each position is convolved with the simulated PSF to obtain an initial estimate of its (ballistic) footprint on the LFM camera. From each footprint, we generate a Boolean mask mi that is one at every pixel behind every microlens that receives a contribution from the ballistic footprint. We collect the set of neuron footprints into a non-negative p × n matrix S_0_, with n being the number of neurons found in the segmentation, and p the number of camera pixels. Also, let Y be the p × t non-negative data matrix (with t the number of time steps in the recording). We then perform a temporal update step by solving the non-negative least squares problem:

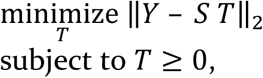

where T is a non-negative n × t matrix T of temporal components, using an iterative solver (see Supplementary Note 2 for convergence criteria). The background components found in the rank-1 matrix factorization performed earlier are inserted as an additional row and column of the S and T matrices, respectively, and therefore updated together with the neuron candidates (Supplementary Note 2).

Next, we perform a spatial update step: We find all sets O^k^ of spatially overlapping components s_i_. For each of these k groups, we form matrices T^k^ that contain all columns ti of T that correspond to spatial components in O^k^, and data matrices Y^k^ that contain only those pixels that fall into the nonzero areas of masks m_i_ in O^k^ (). For each k, we then solve the following non-negative, spatially constrained least-squares problem:

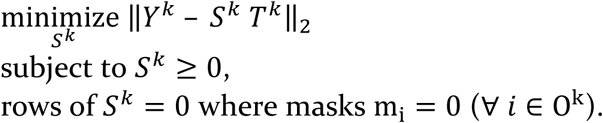

We then iterate the temporal and spatial update steps until convergence (Supplementary Note 2).

Finally, we compute the integral of every spatial component, normalize it to one and scale the temporal component by the integral. We scale the temporal components individually to the standard deviation of the noise they contain (defined as the residual of a Savitzky-Golay fit).

### *In-vivo* Ca^2+^ imaging of head-fixed zebrafish larvae

For zebrafish experiments, elavl3:H2B-GCaMP6s fish (n=4) were imaged 5-8 days post fertilization. This line expresses a nuclear confined Calcium indicator pan-neuronally in a mitfa-/-, roy-/- background. Larvae were immobilized by embedding them in 2% low melting point agarose. For spinal cord recordings, larvae were paralyzed by injection of α-bungarotoxin (125 μM) into the heart cavity at least one hour before the experiment.

### Animal surgery and *in-vivo* Ca^2+^ imaging of awake mice

Surgery and experimental procedures fulfilled the Austrian and European regulations for animal experiments (Austrian §26 Tierversuchsgesetz 2012 – TVG 2012) and were approved by the IACUC of The Rockefeller University. Adult (P90+) male and female C57Bl/6J wild-type mice (n = 10) were anesthetized with isoflurane (2-3% flow rate of 0.5- 0.7 l/min) and placed in a stereotaxic frame (RWD Life Science Co., Ltd. China). After removing the scalp and clearing the skull of connective tissues, a custom-made lightweight metal head-bar was fixed onto the skull with cyanoacrylate adhesive (Krazy Glue) and covered with black dental cement (Ortho-Jet, Lang Dental, USA or Paladur, Heraeus Kulzer, GmbH, Germany). The head-bar was stabilized by anchoring it with up to 3 headless M1.4 screws inserted at the occipital and parietal bones. A circular craniotomy (3-5 mm diameter) was then performed above the imaging site (posterior parietal cortex, PPC, centered at ~2.5 mm caudal and ~1.8 mm lateral; primary motor cortex, M1, -2.5 mm anterior and 1.5 mm lateral; dorsal hippocampus 2.0-2.5 mm caudal and 1.4-1.8 mm lateral to Bregma). With the skull opened and the dura intact, the GECI-carrying virus AAV8:hSyn-GCaMP6m was injected at 4-12 sites (25 nl each, at 10 nl/min; titer ~10^12^ viral particles/ml) with a 400 μm spacing forming a grid near the center of the craniotomy, at a depth of 400-450 μm below dura for PPC and 1200 μm for hippocampus. For data shown in Supplementary Fig. 10, the construct AAV2/1: Hsyn-JRGECO was injected. After the injections, a glass cranial window consisting of a 3-5 mm diameter, #1 thickness (0.16 mm) coverslip was implanted in the craniotomy, flushed with saline solution, placed in contact with the brain surface, and sealed in place using tissue adhesive (Vetbond). The exposed skull surrounding the cranial window was covered with dental cement to build a small chamber for imaging with a water-immersion objective. To access the dorsal hippocampus, a cranial window was implanted after cortical aspiration as previously reported^35,36^. To prevent post-surgical infections and post-surgical pain, the animals were supplied with water containing the antibiotic enrofloxacin (50 mg/Kg) and the pain killer carprofen (5 mg/Kg) for a period of ~7 days. After surgery, animals were returned to their home cages for 2-3 weeks for recovery and viral gene expression before subjecting to imaging experiments. Extreme care was taken to ensure that the dura experienced no damage or major bleeding before and after cranial window implantation. Mice with damaged dura or unclear windows were euthanized and not used for imaging experiments. During imaging sessions, the animals were head-fixed using a customized mount complemented with a head bar holder and a mouse body stabilizer (body jacket) and could freely run on a disk (200 mm diameter). Spontaneous activity was recorded. This considerably reduced animal induced motion of the brain during imaging. A ventilation mask was placed in front of the mouse nose to provide air puff mechanical stimulation to the mouse whiskers and face as well as to provide gas anesthesia on demand. Typical imaging session lasted continuously for 2-10 min.

### Data availability

The data that support the findings of this study are available from the corresponding author upon request.

### Code availability

The custom-written Matlab code implementing Seeded Iterative Demixing is available as Supplementary Software published with the online version of this article under the license terms included with the Supplementary Software package.

